# Comprehension of Antimicrobial Peptides Modulation of the Type VI Secretion System in *Vibrio cholerae*

**DOI:** 10.1101/2022.11.18.517169

**Authors:** Annabelle Mathieu-Denoncourt, Marylise Duperthuy

**Author notes:** Corresponding author: Marylise Duperthuy, Université de Montréal, C.P. 6128, succursale Centre-ville, Montréal, QC, H3C 3J7.

## Abstract

The Type VI secretion System (T6SS) is a versatile weapon used by bacteria for virulence, resistance to grazing and competition with other bacteria. We previously demonstrated that the role of the T6SS in interbacterial competition and in resistance to grazing is enhanced in *Vibrio cholerae* in the presence of subinhibitory concentrations of polymyxin B (PmB). In this study, we performed a global quantitative proteomic analysis by liquid chromatography coupled to mass spectrometry and a transcriptomic analysis by quantitative PCR of the T6SS known regulators in *V. cholerae* grown with and without PmB. The proteome of *V. cholerae* is greatly modified in the presence of PmB at subinhibitory concentrations with more than 39 % of the identified cellular proteins displaying a difference in their abundance, including T6SS-related proteins (Hcp, VasC, TsaB and ClpV). We identified a regulator whose abundance and expression are increased in the presence of PmB, *vxrB*, the response regulator of the two-component system VxrAB. In a *vxrAB* deficient mutant, the expression of *hcp* measured by quantitative PCR, although globally reduced, was not modified in the presence of PmB, confirming its role in *hcp* upregulation with PmB. The upregulation of the T6SS in the presence of PmB appears to be, at least in part, due to the two-component system VxrAB.

**Importance:** The type VI secretion system is important for bacterial competition, virulence and resistance to grazing by predators. In this study, we investigated the regulation leading to the type VI secretion system activation in the presence of polymyxin B (PmB), an antimicrobial used in veterinary and human health to treat infection caused by multi-resistant Gram-negative bacteria, in *V. cholerae*. In addition to making an overall portrait of the modifications to the proteome, we identified the VxrAB two-component system as the main regulator responsible for this activation. Our results provide evidence that subinhibitory concentrations of antimicrobials are responsible for important modifications of the proteome of pathogenic bacteria, inducing the production of proteins involved in virulence, host colonisation, resistance and environmental survival.

## Introduction

*Vibrio cholerae* is a ubiquitous Gram-negative bacterium found in brackish rivers, coastal water and estuaries, or associated with fishes, shellfishes and zooplankton (1). There are more than 200 serotypes of *V. cholerae*, the O1 and O139 causing the cholera disease. The O1 El Tor biotype is responsible for the 7^th^ ongoing pandemic (2). Cholera is acquired by the consumption of contaminated water or food and is characterized by an acute diarrhea, leading to dehydration and eventually to death if not treated (1). *V. cholerae* is endemic in regions where water sanitation facilities are inadequate and access to drinking water is precarious (2, 3). Many virulence and competition effectors allow *V. cholerae* to cause infections, the most important being the cholera toxin and the toxin co-regulated pilus (4, 5).

The type VI secretion system (T6SS), found in more than 25% of Gram-negative bacteria, has been discovered a decade ago in *V. cholerae* (6). The T6SS is a molecular syringe analogous to T4 bacteriophage. It allows the translocation of various toxic effectors into target neighbouring cells. A contraction event of the VipA/B sheath propels a hemolysin-coregulated protein (Hcp) nanotube that punctures the target cell’s envelope (7–9). The effectors have many intracellular targets and modes of action in both eukaryotic and prokaryotic cells, such as disruption of the cytoskeleton by actin cross-linking (10, 11), disruption of the cell membrane (12, 13) and peptidoglycan degradation (14, 15). It makes the T6SS an important weapon against competitor bacteria and predators (12, 13, 16, 17). The T6SS genes are divided in multiple gene clusters, *i.e*. a large cluster (VCA0105-VCA0124) that encodes for the major structural proteins, a protease and the internal regulator *vasH* (VCA0117), and at least 2 auxiliary clusters (VCA0017-VCA0022 and VC1415-VC1421), harbouring *hcp*, *vgrG*, adaptor and effector/immunity proteins (6, 12, 18). The expression of the T6SS genes is complex and involves different regulators (reviewed in (7)), including *vasH*. VasH orchestrates, along with the alternative sigma factor *rpoN*, the expression of the large and auxiliary clusters (7, 19). Most studies on the T6SS in *V. cholerae* were done with non-O1/non-O139 environmental strains that constitutively secrete through their T6SS. However, the regulation appears to vary between strains and seems to differ in clinical isolates as the secretion needs to be triggered. In clinical isolates, high osmolarity and high cell density, involving the regulators *oscR* and *hapR* respectively, are both required for secretion, even though the T6SS is constitutively expressed and produced (20, 21). Globally, the T6SS regulation also depends on other environmental signals, such as bile (22), nucleosides levels (23) and chitin (24–27) through the chitin competence pathway. The global regulators LonA and TsrA also play a role in T6SS regulation. Cold shocks lead to the production of the regulator *cspV*, which modulates the expression of many genes involved in cold shock response, biofilm formation and to the up-regulation of the T6SS (28, 29). The T6SS is also up-regulated by cell wall damages through the two-component system (TCS) VxrAB (30). Bacteria use TCS, a signal transduction device, to regulate the expression of diverse stress or virulence factors in response to variations in their environment and throughout infection (31). TCS are generally composed of a membrane-bound sensor histidine kinase (HK) and a response regulator (RR), which are activated through subsequent phosphorylation. In most TCS, the detection of stimuli by HK leads to the autophosphorylation of its histidine residue, then to the transphosphorylation of the aspartate residue of the RR (32). The RR acts afterwards as a transcriptional regulator. TCS are implicated in various processes such as virulence, stress response, environmental adaptation, and quorum sensing, among others (33). In *V. cholerae, vxrA* and *vxrB* are encoded on the conserved *vxrABCDE* (VCA0565 - VCA0569) (34). VxrC appears to have an inhibitory role in biofìlm formation (30). As for VxrDE, their role is still unknown although it has been suggested that VxrDE have a minor role in *vpsL* expression without affecting the biofilm formation (30). While VxrAB are essential for the colonization of infant mouse model, the contribution of VxrCDE is minor (34).

Antimicrobial peptides (AMPs) are ubiquitous, small, and mostly cationic peptides produced by bacteria and host cells for competition, prevention of infections, or to control the microbiota (35). AMPs are produced in the gut by the microbiota and host cells, and are effective against a broad range of microorganisms such as viruses, fungi and bacteria (36). AMPs have many cellular targets, but the most common mechanism of action is an electrostatic interaction with the negative bacterial membrane that leads to pore formation, the loss of cytoplasmic content and eventually, cell death (37). Polymyxin B (PmB), a cationic AMP produced by *Paenibacillus polymyxa*, binds to the lipid A portion of the LPS at the outer membrane (38). Polymyxins are commonly used in prevention of infections in veterinary medicine and in the treatment of multidrug resistant infections in human health (39). Since the absorption of polymyxins is low, a high percentage is excreted through urine and can accumulate in the environment, including soil and water (40). Polymyxins can thus be found in subinhibitory concentrations in water (41, 42). Subinhibitory concentrations of AMPs are known to have a modulatory effect on virulence, persistence and resistance factors in Gram-negative bacteria (43, 44), including on *V. cholerae* (45–48). El Tor strains are resistant to PmB conversely to the classical strains they replaced (38).

We previously demonstrated that subinhibitory concentrations of PmB increase the expression of Hcp, the structural component of the T6SS syringe in a pandemic strain of *V. cholerae* (46). PmB alone was not sufficient to induce the secretion through the T6SS in an O1 El Tor strain in low osmolarity conditions, but increased its production and secretion in a dose dependant manner. Although it did not increase the resistance of *V. cholerae* to antimicrobial peptides or antibiotics, the increased secretion through the T6SS due to the PmB led to a more efficient elimination of the competitor bacteria *E. coli* and an increased cytotoxicity towards amoebas in a T6SS-dependant manner. Furthermore, the increased expression of both *hcp* genes lets us believe that PmB could act as an activating signal for T6SS production and secretion (44, 46). In this study, we aim to identify the regulatory pathways involved in the over expression and over secretion of Hcp in the presence of subinhibitory concentrations of PmB. A proteomic approach of the cell fraction coupled with a transcriptomic approach using quantitative PCR, allowed to determine that the presence of subinhibitory concentrations of PmB has a complex impact on the proteome of *V. cholerae* and greatly modulates the production of proteins implicated in cellular and metabolic processes, regulation and binding and catalytic activities. Among the proteins that were more abundant in the presence of PmB, we identified VxrB, part of the VxrAB TCS and involved in the T6SS regulation. We further constructed a deficient mutant of *vxrAB* and its Hcp production and expression (*hcp*) were assessed by western blot and quantitative PCR. Our results suggest that *vxrAB* is, at least partly, responsible for the increased expression and secretion of Hcp in the presence of PmB in *V. cholerae*.

## Material & methods

### Strains used in this study

*Vibrio cholerae* O1 El Tor strain A1552, an Inaba clinical strain isolated in 1992 from a Peruvian tourist, as well as its T6SS defective isogenic mutant A1552Δ*hcp1-2* were used for this study (20, 49). Bacterial strains obtained by mutagenesis by natural competence as previously described (see below) (50) and plasmids used in this study are listed in Table I. *V. cholerae* strains were grown on LB agar plates at 37°C. Isolated colonies were inoculated in 5 mL LB broth and incubated 16 h at 37°C with agitation. To obtain the final cultures, 100 μL of the bacterial suspension were transferred into 5 mL of LB broth with 2% NaCl (LB-2%NaCl), with or without 3 μg/mL of PmB (Sigma-Aldrich) and incubated at 37°C with agitation until they reached an optical density at 600nm (OD_600nm_) of 2. When needed, L-arabinose (0.05 % w/v) or antibiotics (chloramphenicol (2 μg/mL), carbenicillin (50 μg/mL)) were added to the media.

**Table I.** Bacterial strains and plasmids used in this study.

| Strain/plasmid | General characteristics | Reference |
| --- | --- | --- |
| *V. cholerae* |  |  |
| A1552 | Wild type strain, O1 El Tor, pathogenic strain isolated from human cholera infection | (49) |
| A1552Δ*hcp1-2* | Hemolysin coregulated protein (Hcp) expression-deficient strain derived from A1552. Deletion of *hcp* genes from auxiliary clusters 1 and 2. Lacks functional T6SS. | (20) |
| A1552Δ*vxrAB*::CmR | A1552 derived strain in which *vxrAB* (VCA0565-66) have been replaced by a chloramphenicol resistance cassette, with promoter and terminator, from pCas9 CR4 | This study |
| A1552Δ*cspV*::CmR | A1552 derived strain in which *cspV* (VCA0933) have been replaced by a chloramphenicol resistance cassette, with promoter and terminator, from pCas9 CR4 | This study |
| *E. coli* |  |  |
| DH5α | F- ϕ80*lacZ*ΔM15 $\Delta$(*lacZYA-argF*) U169 *recA1 endA1 hsdR17*(r<sub>K</sub><sup>-</sup>, m<sub>K</sub><sup>+</sup>) *phoA supE44 λ-thi-1 gyrA96 relA1* | (78) |
| **Plasmids** |  |  |
| pBAD24 | Expression vector. Arabinose inducible promoter, resistance cassette to carbenicillin (https://www.addgene.org/vector-database/1845/) | (65) |
| pBAD24-*vxrAB* | pBAD24 vector with complete *vxrAB* open reading frame from A1552 under *ara* promoter | This study |
| pBAD24-*cspV* | pBAD24 vector with complete *cspV* open reading frame from A1552 under *ara* promoter | This study |
| pCas9 CR4 | Cas9 nuclease under control of pTet promoter with *ssrA* tag and constitutive *tetR* (https://www.addgene.org/62655/) | (52) |

### Proteomic analysis

Proteomic analysis of the bacterial cells grown with or without PmB (3 μg/mL) was performed by liquid chromatography coupled with tandem mass spectrometry, as described before (46). The cellular fraction of *V. cholerae* A1552 cultures at OD_600nm_ of 2 in LB2%NaCl in the presence or in absence of PmB were analyzed. The experiment was conducted in two biological replicates. The proteins were identified using the genome of *V. cholerae* N16961, a strain very closely related to A1552. The data were analyzed using the Scaffold V.5 software (protein threshold: 99%, with at least two peptides identified and a false discovery rate of 1%) and we removed the proteins identified in only one of the replicates. The proteins with differential abundance were then subjected to a Gene Ontology (GO) term enrichment (51).

### Mutant construction and complementation

A1552Δ*vxrAB*::CmR and A1552Δ*cspV*::CmR defective mutants were obtained using the optimized natural competence protocol by Marvig & Blokesch (50). All the PCR amplicons were obtained with Thermo Fisher^™^ oligonucleotide primers listed in Table II, using Taq DNA Polymerase with Standard buffer from New England Biolabs^®^, according to the manufacturer’s instructions. The chloramphenicol resistance cassette (CmR), with its promoter and terminator from pCas9 CR4 (52) (Table I), was amplified by PCR. Up and downstream regions of target genes of about 1000 bp were amplified from A1552 genomic DNA by PCR using primers adding 15 bp homology to CmR 5’ and 3’ extremities (Table II). The upstream region, the CmR cassette and the downstream region were bound together by overlap PCR and purified using Monarch DNA Gel extraction kit (New England Biolabs^®^).

**Table II.**
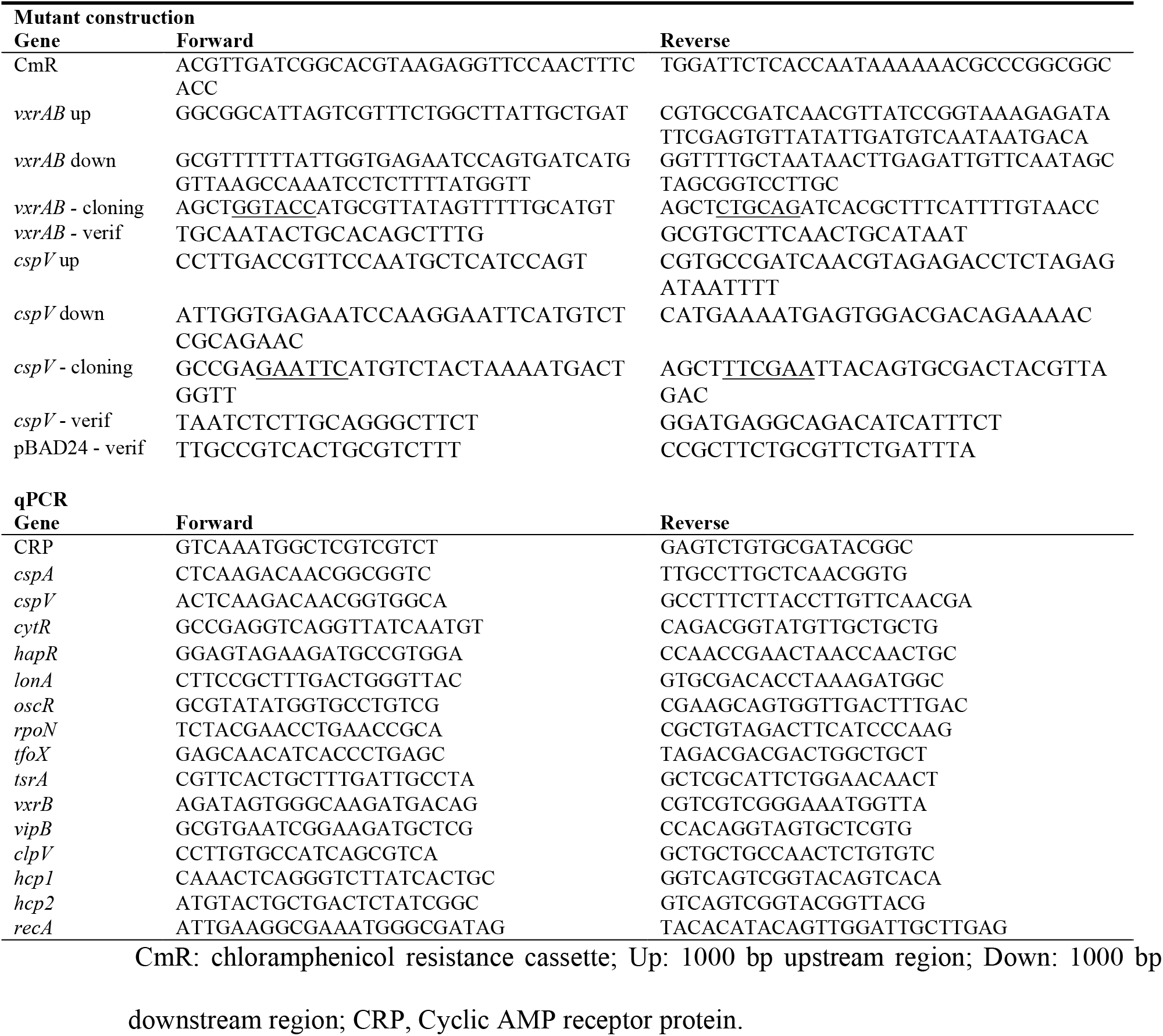
Primers used in this study.

For the natural transformation, briefly, *V. cholerae* wild type strain A1552 was grown over night in LB at 30°C with agitation. The culture was diluted 1:50 in fresh LB and incubated at 30°C until it reached an OD_600nm_ of 0.5. Bacteria were washed by centrifugation at 5500 x *g* for 5 min and suspended in M9 minimal media (Sigma) supplemented with 32 mL of MgSO_4_ 1 M and 5.1 mL of CaCl_2_ 1 M per liter (50). The bacteria were diluted 1:2 in fresh M9 media. Chitin powder (Sigma-Aldrich) was added, and the culture was grown overnight at 30°C with agitation. Two hundred ng of purified PCR amplicon were added to the culture, and the bacteria were further incubated 24 h at 30°C. To detach the bacteria from chitin, fresh M9 medium was added to the culture and the bacteria were vortexed vigorously for 2 min. The chitin was pelleted by quick centrifugation and the supernatant containing free bacteria was centrifuged at 5000 x *g* for 5 min. The bacteria were suspended in 500 μL of PBS and plated on LB agar supplemented with 2 μg/mL chloramphenicol (Fisher). Plates were incubated at 30°C for 3 days. Colonies grown on agar supplemented with chloramphenicol were further isolated on agar plates with chloramphenicol to confirm the phenotype. Deficient mutants were confirmed with PCR using either *cspV* – verif or *vxrAB* – verif primers (Table II) and sequencing of the insertion region.

For complementation, the complete open reading frame of *cspV* or *vxrAB* was amplified by PCR from purified A1552 genomic DNA using the Thermo Fisher^™^ oligonucleotide primers adding restriction sites listed in Table II. The amplicons and pBAD24 vector (Table I) were digested using HindIII and EcoRI (*cspV*) or PstI and KpnI (*vxrAB*) from New England Biolabs^®^ according to the manufacturer’s instructions. Plasmids and amplicons were purified from agarose gel using Monarch Gel purification kit (New England Biolabs^®^) and ligated using T4 ligase from New England Biolabs^®^. The final construction was introduced by heat shock into thermocompetent *E. coli* DH5α at 42°C for 30 s (53). The bacteria were then suspended in fresh LB and incubated 1.5 h at 37°C. Bacteria were selected on LB agar supplemented with carbenicillin (50 μg/mL) (Fisher). Plasmids were extracted from *E. coli* using PureYield^™^ Plasmid Miniprep System (Promega) and suspended in milliQ water (Thermo Fisher).

*V. cholerae* was grown in LB for 16 h at 37°C and were made electrocompetent by successive washes in sterile water supplemented with 10% glycerol. The constructions were electroporated in the corresponding strains at 1.275 kV, 25 Ω in 1 mm electroporation cuvettes (Thermo Fisher). The bacteria were incubated 1.5 h at 37°C in LB, then plated on LB agar containing 50 μg/mL carbenicillin. Colonies were screened by PCR using pBAD24 – verif primers (Table II) to verify the construction.

### Growth curves

*V. cholerae* was grown 16 h at 37 °C with agitation in LB, with carbenicillin when needed. A 1:50 dilution in fresh LB was done, and the bacteria were grown at 37°C to an OD_600nm_ of 0.2. Then, 0.05 % L-arabinose was added, and the bacteria were grown until they reached OD_600nm_ = 0.3. They were further diluted 1:50 in LB2%NaCl - 0.05 % L-arabinose, with or without 3 μg/mL of PmB, in 50 ml Falcon tubes, or 1:3000 in 96 wells plates. The bacterial growth was followed by reading the OD_600nm_ every 15 min, at 37°C with agitation. Data were obtained from at least three independent experiments, in technical triplicates.

### RNA extraction, cDNA synthesis and quantitative PCR (RT-qPCR)

*V. cholerae* was grown to an OD_600nm_ of 0.5 at 37°C in 10 mL of LB2%NaCl, with or without 3 μg/mL of PmB. The bacteria were suspended in 1 mL TRIzol solution (Invitrogen) and the total RNA was extracted according to the manufacturer’s instructions. Five hundred nanograms of RNA were retrotranscribed to cDNA using QuantiNova Reverse Transcription Kit (Qiagen). RNA and cDNA purity and quality were assessed by nanodrop and migration on 2% agarose gel, respectively. Quantitative PCR analysis was done as described before (46) with primers listed in Table II. The relative *hcp* and regulators’ expression was calculated in PmB treated bacteria in comparison to non-treated cells using QuantStudio^™^ Desing and Analysis Software (Thermo Fisher) v1.5.1 and normalized using *recA*. The results were obtained from at least 3 independent experiments, in technical triplicates.

### Western blot

Bacteria were grown to an OD_600nm_ of 2 in LB2%NaCl, a condition that activates the T6SS secretion in *V. cholerae* A1552, supplemented with - 0.05 % L-arabinose and 3 μg/mL of PmB. Samples were treated and western blot conducted as described before (46). Results are representative of at least three independent experiments.

### Statistical Analysis

All data are expressed as mean ± SD and were analyzed for significance using the SigmaPlot (version). Student’s *t*-tests were used to compare conditions between 2 groups. Single way ANOVA was used for multiple groups comparison. A result was considered as significant when *p* value < 0.05 (*).

## Results

### Proteomic analysis of *V. cholerae’s* cells grown with subinhibitory concentrations of PmB

To identify the cellular proteins whose abundance is modified in the presence of PmB, a quantitative proteomic analysis by liquid chromatography coupled to mass spectrometry was performed on *V. cholerae* A1552 bacterial cells grown in LB2%NaCl with or without 3 μg/ml of PmB, a concentration that does not affect bacterial growth but significantly increases the expression and secretion of Hcp (46). Our proteomic analysis identified a total of 22,819 peptides corresponding to 473 proteins of 7 to 184 kDa. After data curation, 454 proteins were obtained. We calculated the relative abundance of each protein in the presence of PmB in comparison with the control without PmB and determined that 177 proteins (39%) showed a modified abundance in the presence of PmB (Tables IV to VII). Among them, 130 were more abundant in the presence of PmB (Table IV), 17 were less abundant in its presence (Table V), 26 were only found in its presence (Table VI), and 4 were found only in its absence (Table VII). The proteins showing no modulation are presented in Table SI. The correlation of the biological replicates was analyzed using scatter plots and demonstrated a good correlation with R-squared of more than 0.9 (Figure 1AB).

**Figure 1.**
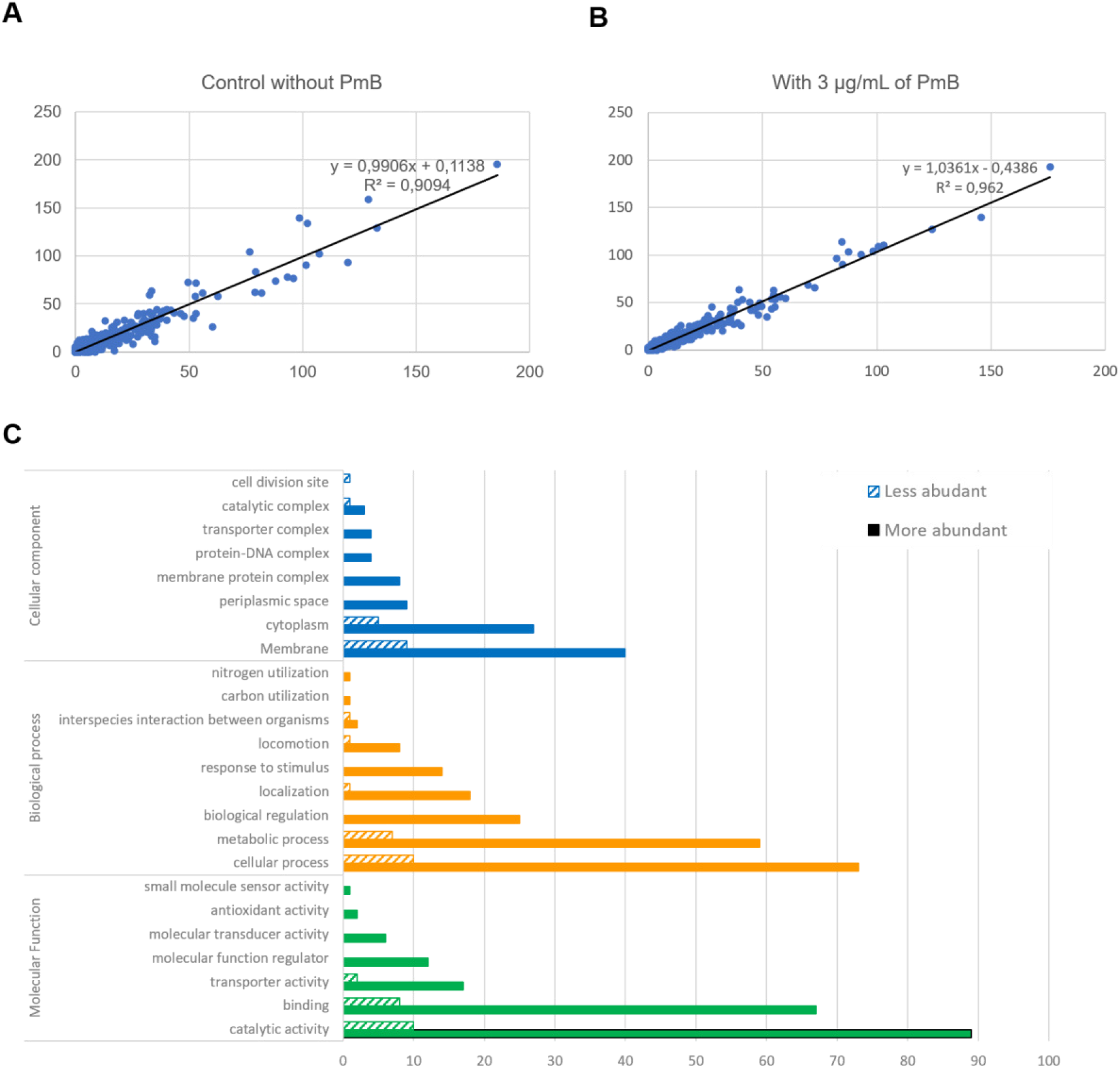
Polymyxin B modifies the proteome of *V. cholerae*. A quantitative proteomic analysis by liquid chromatography coupled to mass spectrometry of *V. cholerae* A1552 grown in Type VI Secretion System activating conditions, with or without 3 μg/ml of Polymyxin B (PmB) was performed. Scatter plot analysis using the total number of spectra per protein was performed to determine the correlation between the biological duplicates of the sample without PmB **(A)** and with 3 μg/ml of PmB **(B)**. A GoAnnotation analysis of the proteins identified determined the molecular functions, implication in biological processes and localization of the proteins **(C)**.

**Table III.** Known regulators of the Type VI secretion system of *V. cholerae* identified in our proteomic analysis.

| Gene |  | Found in Analysis | Modified abundance in the presence of PmB | Reference |
| --- | --- | --- | --- | --- |
| VC_1021 | LuxO | No | NA | (20) |
| VC_2677 | CytR | Yes | No | (23) |
| VC_1153 | TfoX | No | NA | (24, 25, 27) |
| VC_2614 | CRP | Yes | No | (20) |
| VC_0583 | HapR | No | NA | (20, 25) |
| VC_0396 | QstR | No | NA | (24) |
| VC_1722 | TfoY | Yes | No | (27) |
| VC_0117 | VasH | No | NA | (6) |
| VC_2529 | RpoN | Yes | No | (6) |
| VC_0070 | TsrA | Yes | No | (76) |
| VC_A0566 | VxrB | Yes | Increased | (34) |
| VC_A0933 | CspV | No | NA | (28) |
| VC_A0029 | OscR | Yes | No | (21) |
| VC_1920 | LonA | No | NA | (79, 80) |

**Table IV.**
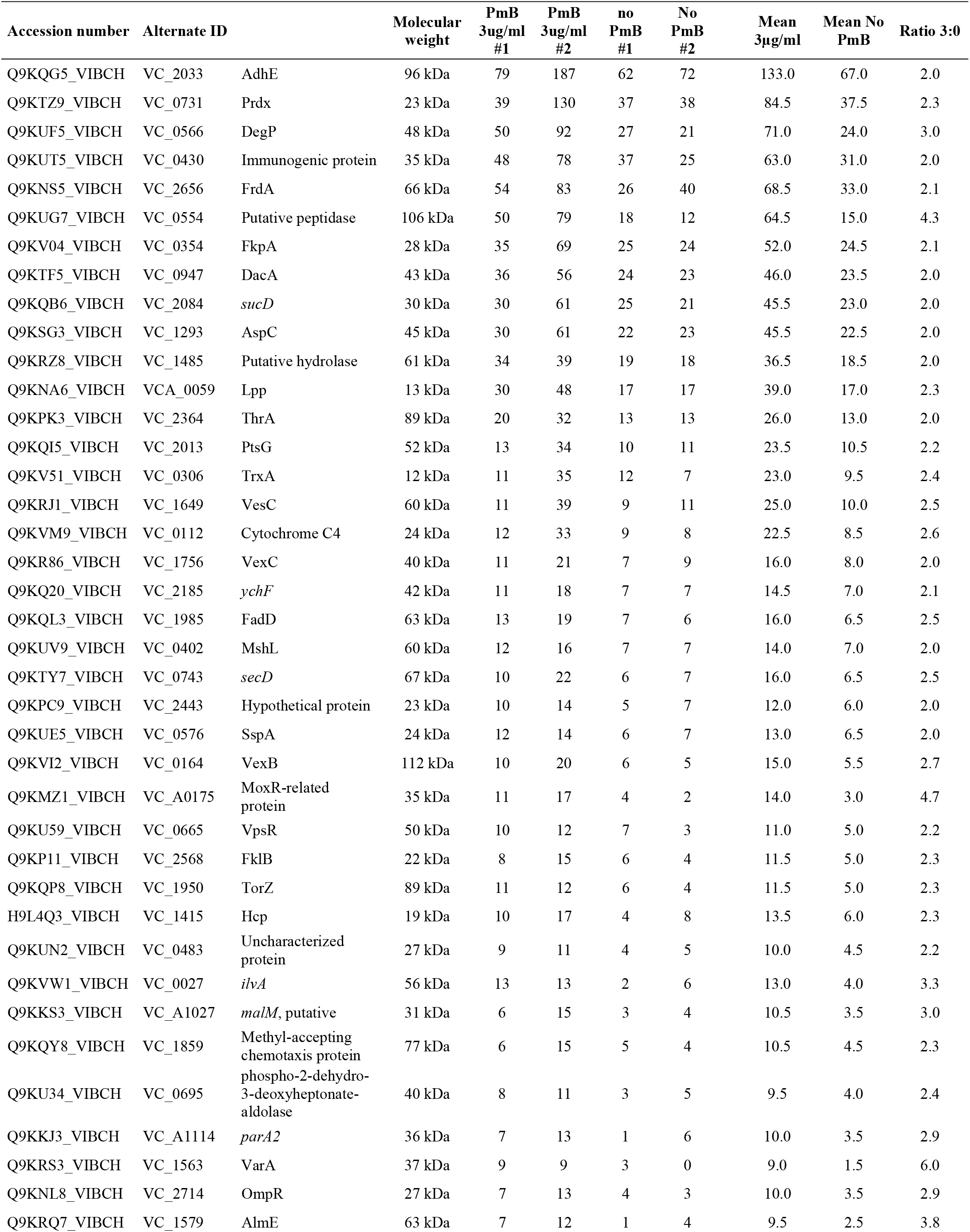
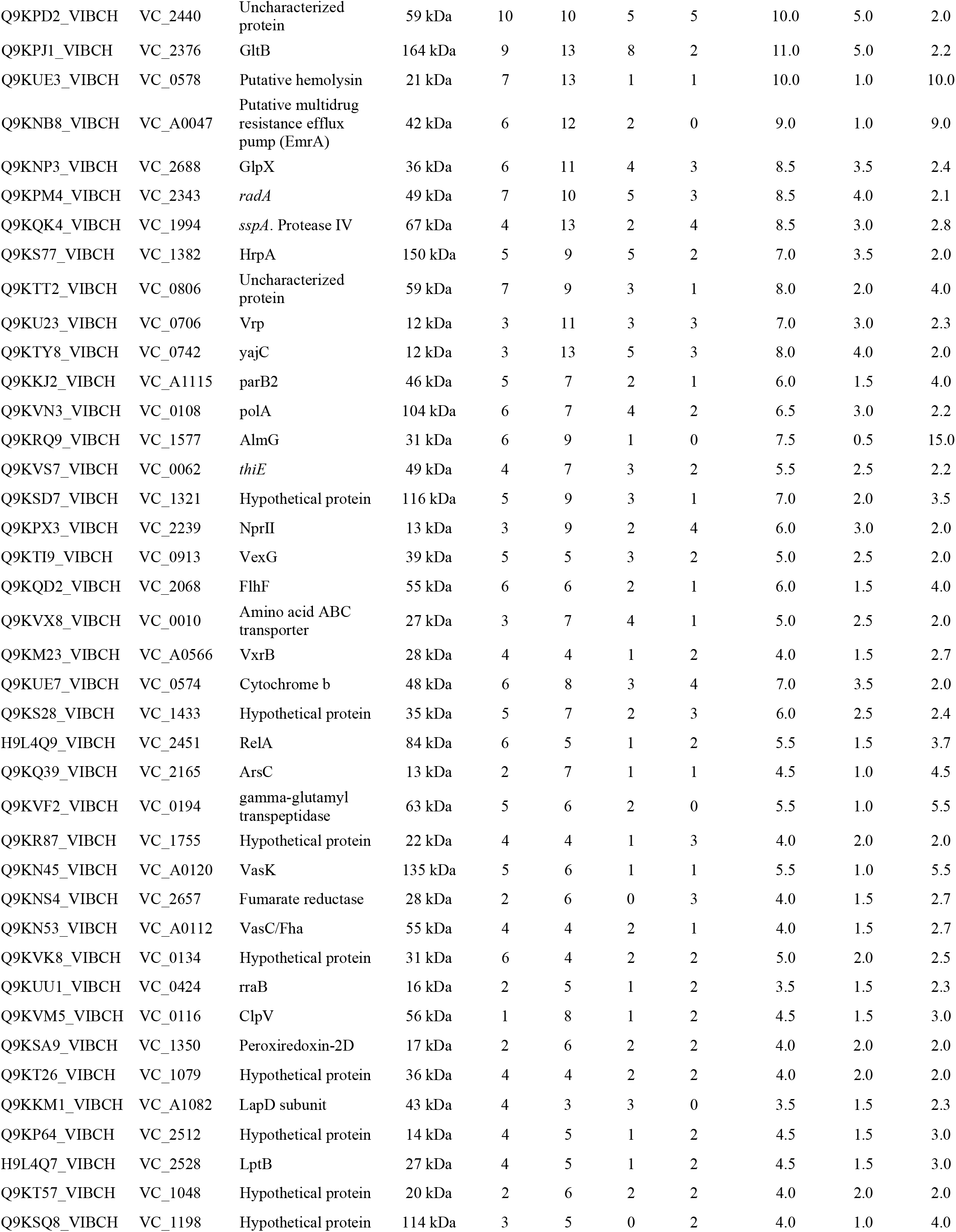
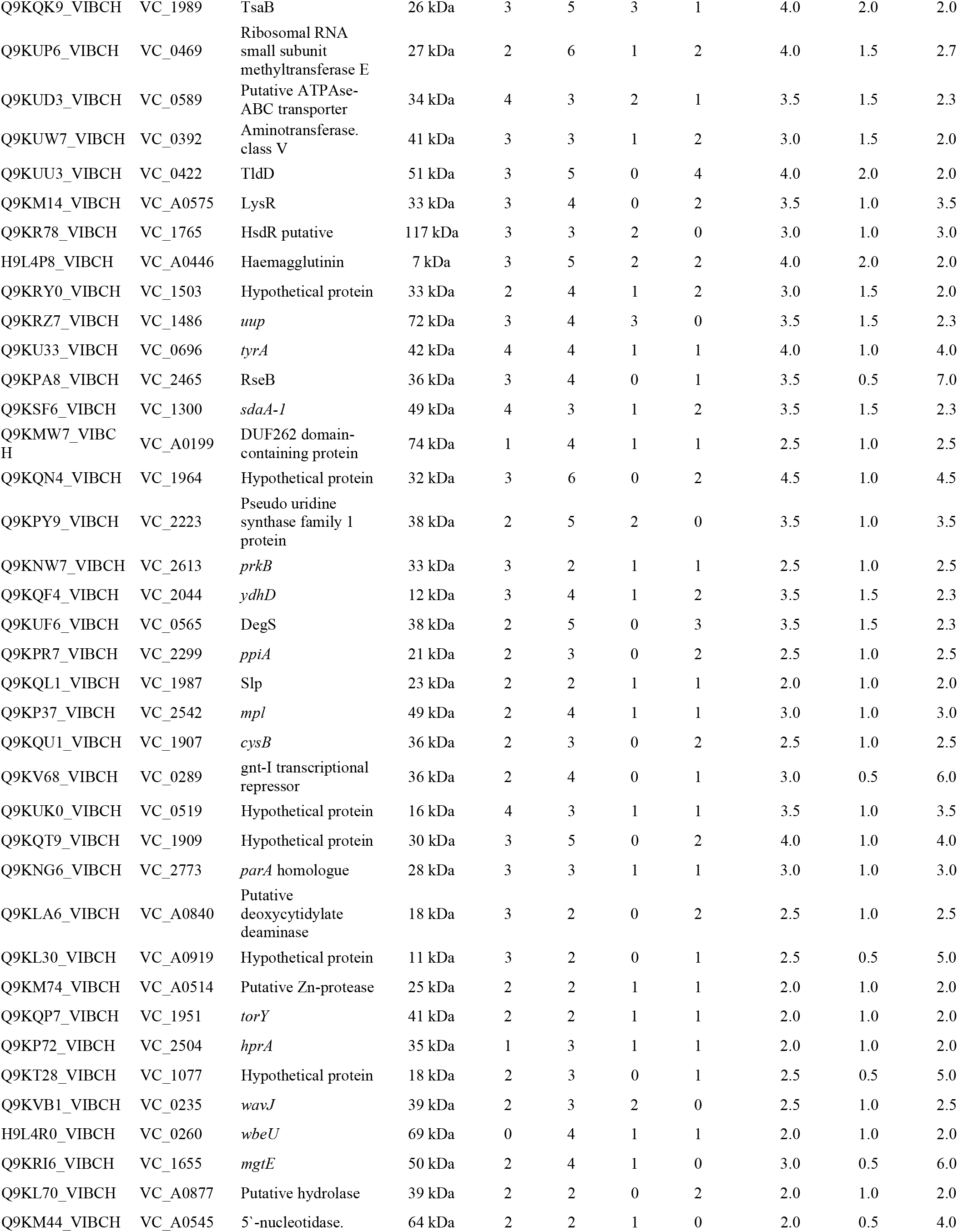
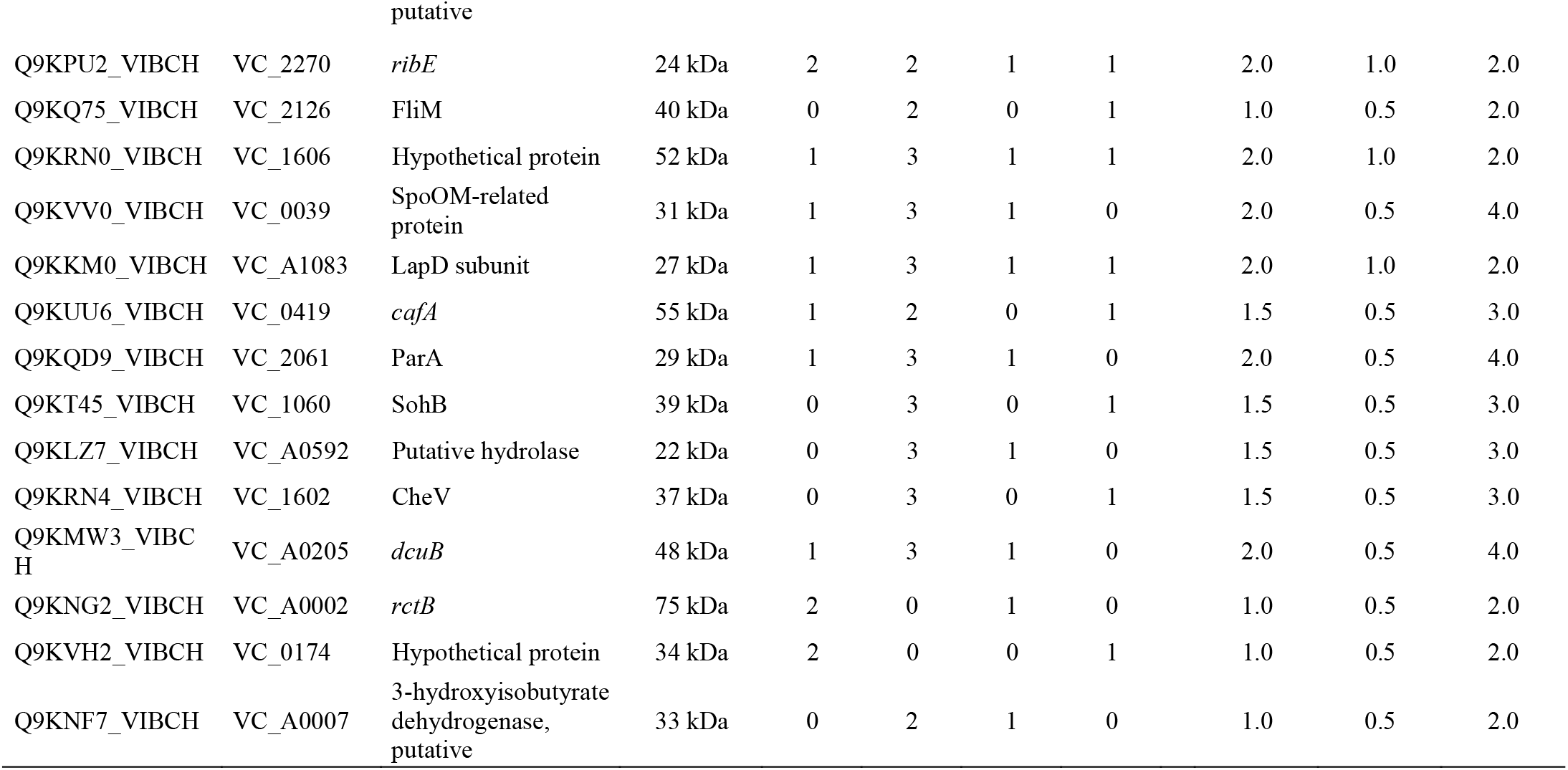
Cellular proteins that are more abundant in the presence of polymyxin B in *V. cholerae* A1552.

**Table V.** Cellular proteins that are less abundant in the presence of polymyxin B in *V.cholerae* A1552.

| Accession number | Alternate ID |  | Molecular weight | PmB 3ug/ml #1 | PmB 3ug/ml #2 | no PmB #1 | No PmB #2 | Mean 3µg/ml | Mean No PmB | Ratio 3:0 |
| --- | --- | --- | --- | --- | --- | --- | --- | --- | --- | --- |
| Q9KQZ3_VIBCH | VC_1854 | *ompT* | 40 kDa | 3 | 7 | 25 | 26 | 5.0 | 25.5 | 0.2 |
| Q9KRP6_VIBCH | VC_1590 | AlsS | 62 kDa | 2 | 2 | 4 | 4 | 2.0 | 4.0 | 0.5 |
| Q9KR21_VIBCH | VC_1826 | IIABC mannose permease | 66 kDa | 1 | 3 | 2 | 8 | 2.0 | 5.0 | 0.4 |
| Q9KM21_VIBCH | VC_A0568 | VxrD | 42 kDa | 0 | 2 | 5 | 5 | 1.0 | 5.0 | 0.2 |
| Q9KS84_VIBCH | VC_1375 | Hypothetical protein | 27 kDa | 0 | 1 | 3 | 1 | 0.5 | 2.0 | 0.3 |
| Q9KL64_VIBCH | VC_A0883 | *makA* | 39 kDa | 1 | 0 | 4 | 4 | 0.5 | 4.0 | 0.1 |
| Q9KLG5_VIBCH | VC_A0779 | Fad-dependant oxidoreductase | 53 kDa | 0 | 1 | 3 | 2 | 0.5 | 2.5 | 0.2 |
| Q9KT68_VIBCH | VC_1037 | *mrp* | 41 kDa | 1 | 1 | 4 | 0 | 1.0 | 2.0 | 0.5 |
| Q9KQJ0_VIBCH | VC_2008 | Pyruvate kinase II | 52 kDa | 1 | 0 | 2 | 0 | 0.5 | 1.0 | 0.5 |
| Q9KTX2_VIBCH | VC_0758 | Hypothetical protein | 43 kDa | 1 | 0 | 1 | 4 | 0.5 | 2.5 | 0.2 |
| Q9KTA5_VIBCH | VC_0998 | HubP | 178 kDa | 0 | 1 | 2 | 1 | 0.5 | 1.5 | 0.3 |
| Q9KUQ6_VIBCH | VC_0459 | Hypothetical protein | 21 kDa | 1 | 0 | 1 | 1 | 0.5 | 1.0 | 0.5 |
| Q9KUT7_VIBCH | VC_0428 | *uspA* | 16 kDa | 1 | 0 | 1 | 3 | 0.5 | 2.0 | 0.3 |
| Q9KPH0_VIBCH | VC_2398 | *ftsA* | 46 kDa | 1 | 0 | 2 | 2 | 0.5 | 2.0 | 0.3 |
| Q9KT65_VIBCH | VC_1040 | Corrinoid adenosyl transferase | 22 kDa | 0 | 1 | 1 | 1 | 0.5 | 1.0 | 0.5 |
| Q9KMF5_VIBCH | VC_A0405 | hypothetical protein | 23 kDa | 1 | 0 | 2 | 2 | 0.5 | 2.0 | 0.3 |
| E5EUX8_VIBCL | VC_A1025 | *nagB* | 23 kDa | 1 | 0 | 2 | 0 | 0.5 | 1.0 | 0.5 |

**Table VI.** Cellular proteins that are only found in the presence of polymyxin B in *V.cholerae* A1552.

| Accession number | Alternate ID |  | Molecular weight | PmB 3ug/ml #1 | PmB 3ug/ml #2 | No PmB #1 | No PmB #2 | Mean 3µg/ml | Mean No PmB |
| --- | --- | --- | --- | --- | --- | --- | --- | --- | --- |
| Q9KKN3_VIBCH | VC_A1069 | Methyl-accepting chemotaxis protein | 70 kDa | 6 | 13 | 0 | 0 | 9.5 | 0.0 |
| Q9KSD9_VIBCH | VC_1319 | VprB | 50 kDa | 8 | 11 | 0 | 0 | 9.5 | 0.0 |
| Q9KKX6_VIBCH | VC_A0974 | Methyl-accepting chemotaxis protein | 61 kDa | 6 | 8 | 0 | 0 | 7.0 | 0.0 |
| Q9KV74_VIBCH | VC_0282 | Methyl-accepting chemotaxis protein | 61 kDa | 2 | 1 | 0 | 0 | 1.5 | 0.0 |
| Q9KSK4_VIBCH | VC_1252 | CinA-like protein | 46 kDa | 3 | 7 | 0 | 0 | 5.0 | 0.0 |
| Q9KQ67_VIBCH | VC_2135 | FlrC | 52 kDa | 2 | 5 | 0 | 0 | 3.5 | 0.0 |
| Q9KMZ9_VIBCH | VC_A0167 | CalB | 29 kDa | 2 | 5 | 0 | 0 | 3.5 | 0.0 |
| Q9KQ69_VIBCH | VC_2133 | FliF | 64 kDa | 3 | 7 | 0 | 0 | 5.0 | 0.0 |
| Q9KP53_VIBCH | VC_2523 | Hypothetical protein | 35 kDa | 2 | 7 | 0 | 0 | 4.5 | 0.0 |
| Q9KLR3_VIBCH | VC_A0679 | NapB | 19 kDa | 2 | 4 | 0 | 0 | 3.0 | 0.0 |
| Q9KLU7_VIBCH | VC_A0644 | NADH oxidase, putative | 62 kDa | 2 | 5 | 0 | 0 | 3.5 | 0.0 |
| Q9KPF1_VIBCH | VC_2417 | RecJ | 65 kDa | 2 | 4 | 0 | 0 | 3.0 | 0.0 |
| Q9KQ71_VIBCH | VC_2130 | *fliL* | 47 kDa | 2 | 2 | 0 | 0 | 2.0 | 0.0 |
| Q9KNB7_VIBCH | VC_A0048 | Hypothetical protein | 15 kDa | 1 | 3 | 0 | 0 | 2.0 | 0.0 |
| Q9KP12_VIBCH | VC_2567 | LysM domain-containing protein | 22 kDa | 1 | 1 | 0 | 0 | 1.0 | 0.0 |
| Q9KV14_VIBCH | VC_0344 | AmiB | 64 kDa | 1 | 1 | 0 | 0 | 1.0 | 0.0 |
| Q9KTA8_VIBCH | VC_0995 | EIIB/EIIC protein | 55 kDa | 2 | 1 | 0 | 0 | 1.5 | 0.0 |
| Q9KSE9_VIBCH | VC_1308 | TyrR | 58 kDa | 2 | 2 | 0 | 0 | 2.0 | 0.0 |
| Q9KQM5_VIBCH | VC_1973 | *menB* | 33 kDa | 1 | 4 | 0 | 0 | 2.5 | 0.0 |
| Q9KTC9_VIBCH | VC_0973 | Hypothetical protein | 30 kDa | 3 | 0 | 0 | 0 | 1.5 | 0.0 |
| Q9KKM3_VIBCH | VC_A1080 | LapC | 54 kDa | 1 | 2 | 0 | 0 | 1.5 | 0.0 |
| Q9KTD7_VIBCH | VC_0965 | PtsI | 63 kDa | 1 | 1 | 0 | 0 | 1.0 | 0.0 |
| Q9KRR8_VIBCH | VC_1568 | VarF | 25 kDa | 1 | 1 | 0 | 0 | 1.0 | 0.0 |
| Q9KUR3_VIBCH | VC_0452 | MutY | 40 kDa | 1 | 2 | 0 | 0 | 1.5 | 0.0 |
| Q9KT72_VIBCH | VC_1033 | Cation transport ATPase | 81 kDa | 1 | 2 | 0 | 0 | 1.5 | 0.0 |
| Q9KMZ2_VIBCH | VC_A0174 | DUF58 domain-containing protein | 35 kDa | 2 | 1 | 0 | 0 | 1.5 | 0.0 |

**Table VII.**
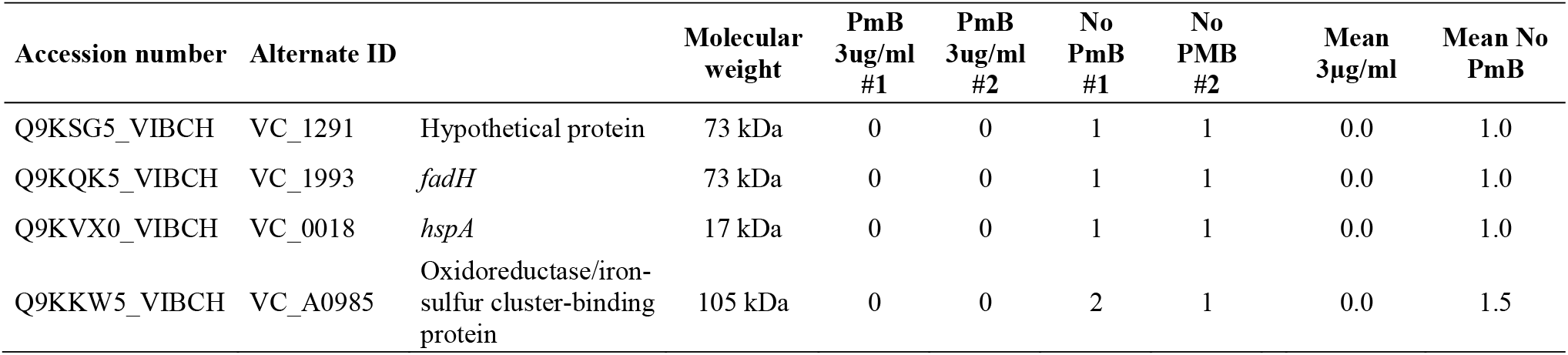
Cellular proteins that are only found in absence of polymyxin B in *V.cholerae* A1552.

An analysis of the proteins that are modulated by PmB using Go Annotation showed that most modulated proteins have catalytic and/or binding function, with 99 and 75 proteins with these functions, respectively (Figure 1C). Regulatory proteins, transporters, transducers and sensor proteins were also modulated by PmB. The analysis showed that the proteins that are more abundant in the presence of PmB were mainly implicated in biological (59) and cellular (75) processes (Figure 1C). Regulation, response to stimulus and locomotion were the biological processes that were greatly modified by PmB.

Among the proteins with an increased abundance in the presence of PmB, many have a role in AMPs or PmB resistance (54). For example, VexB, which is part of the efflux system VexAB (55), was 2.7 times more abundant in the presence of PmB, and AlmG and AlmE, which are directly responsible for PmB resistance by LPS modification (56), were 15 and 3.8 times more abundant with PmB, respectively (Table IV). VprB, only present in the presence of PmB (Table VI), is part of the TCS VprAB that activates the expression of AlmGE (57). The protease DegS was also 2.3 times more abundant in the presence of PmB (Table IV). It is responsible for the activation of the alternate sigma factor RpoE, which activates the repair of AMPs induced damages in the bacterial envelope (58). VarF is part of an antibiotic efflux pump (59) and is only found in the presence of PmB (Table VI). OmpR, the RR to the EnvR-OmpR TCS, is a repressor of virulence factors in response to stress envelope, was also more abundant in the presence of PmB (Table IV) (60).

As for the T6SS related proteins, many showed an increased abundance in the presence of PmB, although a lot of the structural components were not identified in our analysis. As expected, the main component of the T6SS syringe, Hcp (VC_1415, VC_A0017), was also more abundant (2.3 times) in the presence of PmB (46). VasC (VC_A0112), a cytoplasmic protein with an FHA domain essential for secretion (61), and ClpV (VC_A0116), the ATPase responsible for recycling of the contractile sheath, presented abundances that were 2.7 and 3.0 times higher in the presence of PmB, respectively. The abundance of VasK (VC_A0120), a part of the inner-membrane complex (62) was increased by 5.5 times by PmB (Table IV). TsaB (VC_1989), the immunity protein to the VgrG-3 that targets the peptidoglycan, had an increased abundance (2.0 times) in the presence of PmB (Table IV). The abundance of VipA (VC_A0107) and VipB (VC_A0108), the small and large subunits of the contractile sheath, was not modified by PmB (Table S1).

We paid a particular attention to the effect of PmB on the T6SS known regulators. As shown in the table III, our proteomic analysis identified many of them. The abundance of most of these regulators was not significantly affected by the presence of PmB, except for VxrB that was 2.7 times more abundant in the presence of PmB (Table III).

### Transcriptomic analysis of known T6SS regulators in the presence of PmB

To confirm the results obtained in the proteomic analysis and to include regulators that were not identified in this analysis, we performed a transcriptomic analysis of all the known T6SS regulators of *V. cholerae* A1552 grown in LB2%NaCl supplemented or not with 3 μg/ml of PmB. To do so, a quantitative PCR approach was used and the relative expression of known T6SS regulators was calculated by comparing the number of transcripts from bacterial cells grown with and without PmB and normalized with *recA* (Figure 2A). Although statistically significant, the expression of cyclic AMP receptor protein (CRP) (1.23x), *tfoY* (1.15x), *tsrA* (1.17x) and *osrC* (1.26x) was only slightly higher in the presence of PmB, and was not considered for further analysis. Interestingly, the expression of *vxrB* was 2.1 times higher in the presence of PmB than in its absence, while the expression of *cspV* was decreased by 2.43 times in its presence.

**Figure 2.**
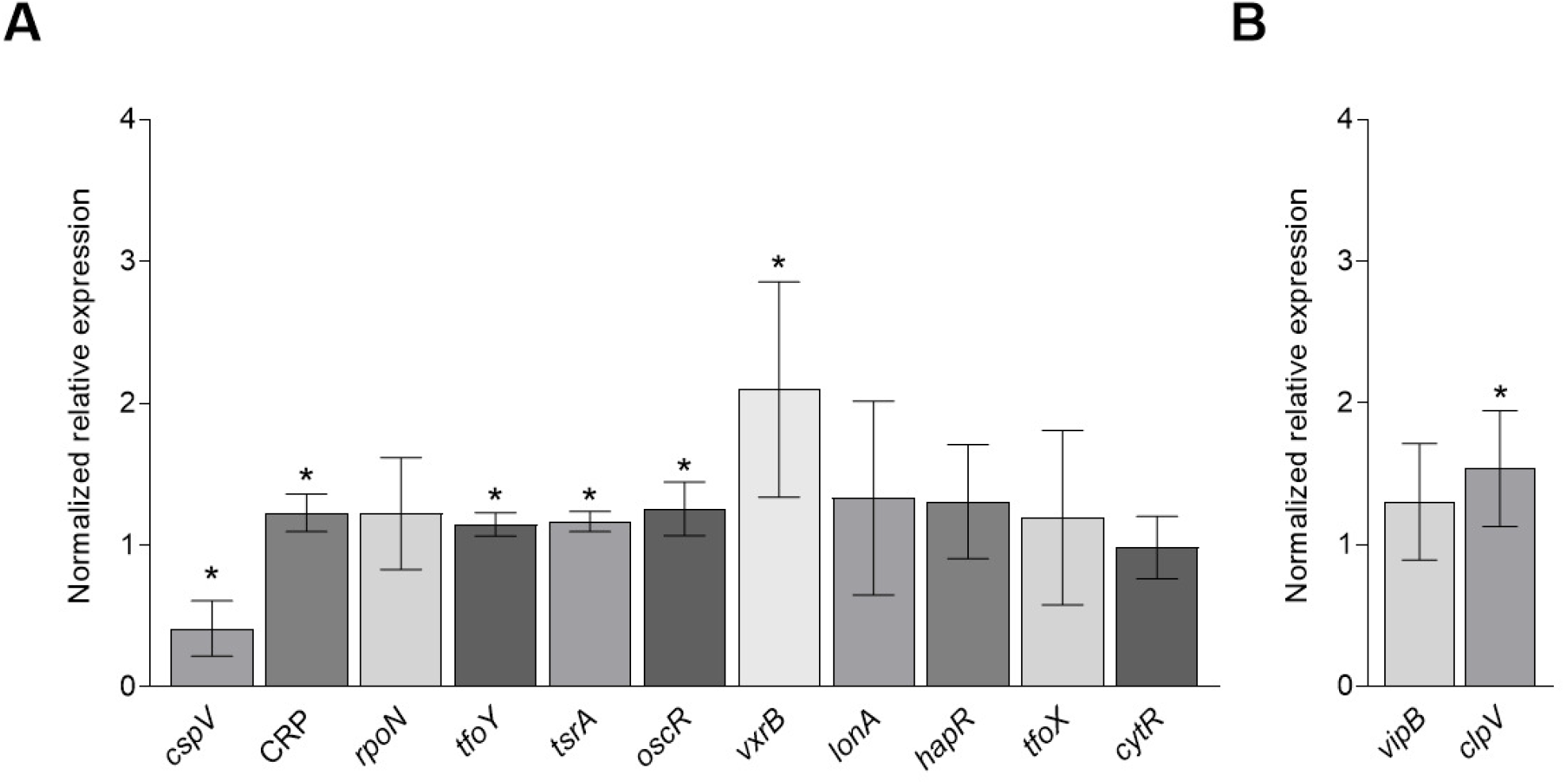
Normalized relative expression of known T6SS regulators (A) and structural components (B) in *V. cholerae* grown with polymyxin B. *V. cholerae* A1552 was grown to midlog phase in LB2%NaCl supplemented or not with 3 μg/ml of polymyxin B (PmB). Total RNA was extracted from cultures and retrotranscribed. The expression of selected regulators and structural components was measured by quantitative PCR in the presence of PmB in comparison to the non-treated cells, and normalized using *recA*. Data represent mean ± SD of three independent experiments conducted in triplicates. Asterisk represents a significant difference in expression between treated and non-treated cells (*P*<0.05). CRP, cyclic AMP receptor protein.

We also assessed the expression of 2 genes encoding structural proteins of the T6SS, the main component of the contractile sheath, VipB, and ClpV, the ATPase protein responsible for the recycling of the sheath after a contraction event (Figure 2B) (63, 64). Our results showed that the expression of *vipB* is not modified by PmB (Figure 2B). This result is in line with our previous work and the proteomic analysis of this study that showed no difference in protein abundance in the bacterial cell (46). The expression of *clpV* is 1.54 times higher in the presence of PmB (Figure 2B), which is also in line with our proteomic analysis.

Based on the results of both proteomic and transcriptomic analysis showing an increased abundance and expression of VxrB and *vxrB*, respectively, in the presence of PmB, and a decreased expression of *cspV* in the presence of PmB, we constructed defective mutants of the regulators *vxrAB* and *cspV* using natural competence (50) and complemented strains using pBAD24 vector (65). The deletion of *vxrAB* (Figure S1A) or *cspV* (Figure S2A) had no major consequence on bacterial growth. The growth was slightly impaired by complementation and PmB for A1552Δ*vxrAB*::CmR (Figure S1B), but not for A1552Δ*cspV*::CmR (Figure S2B). Like A1552, A1552Δ*cspV*::CmR and A1552Δ*vxrAB*::CmR grew in the presence of PmB at a concentration up to 100 ug/ml, although the growth of A1552Δ*vxrAB*::CmR was impaired in all PmB conditions (result not shown), an observation in line with previous studies (66).

### Quantitative PCR analysis of *hcp* shows that *vxrAB*, but not *cspV*, is responsible for its upregulation in the presence of PmB

To determine the effect of the *vxrAB* and *cspV* deletion on the regulation of *hcp* in the presence of PmB, we performed a quantitative PCR analysis. *V. cholerae* A1552, A1552Δ*cspV*::CmR and A1552Δ*vxrAB*::CmR were grown in LB2%NaCl with or without PmB. We quantified the expression of *hcp1* or *hcp2* by quantitative PCR in the presence of PmB, in comparison to the non-treated cells, and normalized using *recA* (Figure 3 and S3). The expression of *hcp1* and *hcp2* in A1552 treated with PmB was 3.03 and 2.16 times higher, respectively, similarly to what we observed before (46) (Figure 3). For A1552Δ*cspV*::CmR, the expression of *hcp1* and *hcp2* is similar to the wildtype strain, and an upregulation of both *hcp1* and *hcp2* is still observed in the presence of PmB (Figure S3)., indicating that, upon *cspV* mutation, the T6SS is activated by PmB. However, upon *vxrAB* mutation, the expression of *hcp1* and *-2* was significantly reduced, as expected (34), but was not increased in the presence of PmB (Figure 3), indicating that the upregulation due to the PmB is no longer effective.

**Figure 3.**
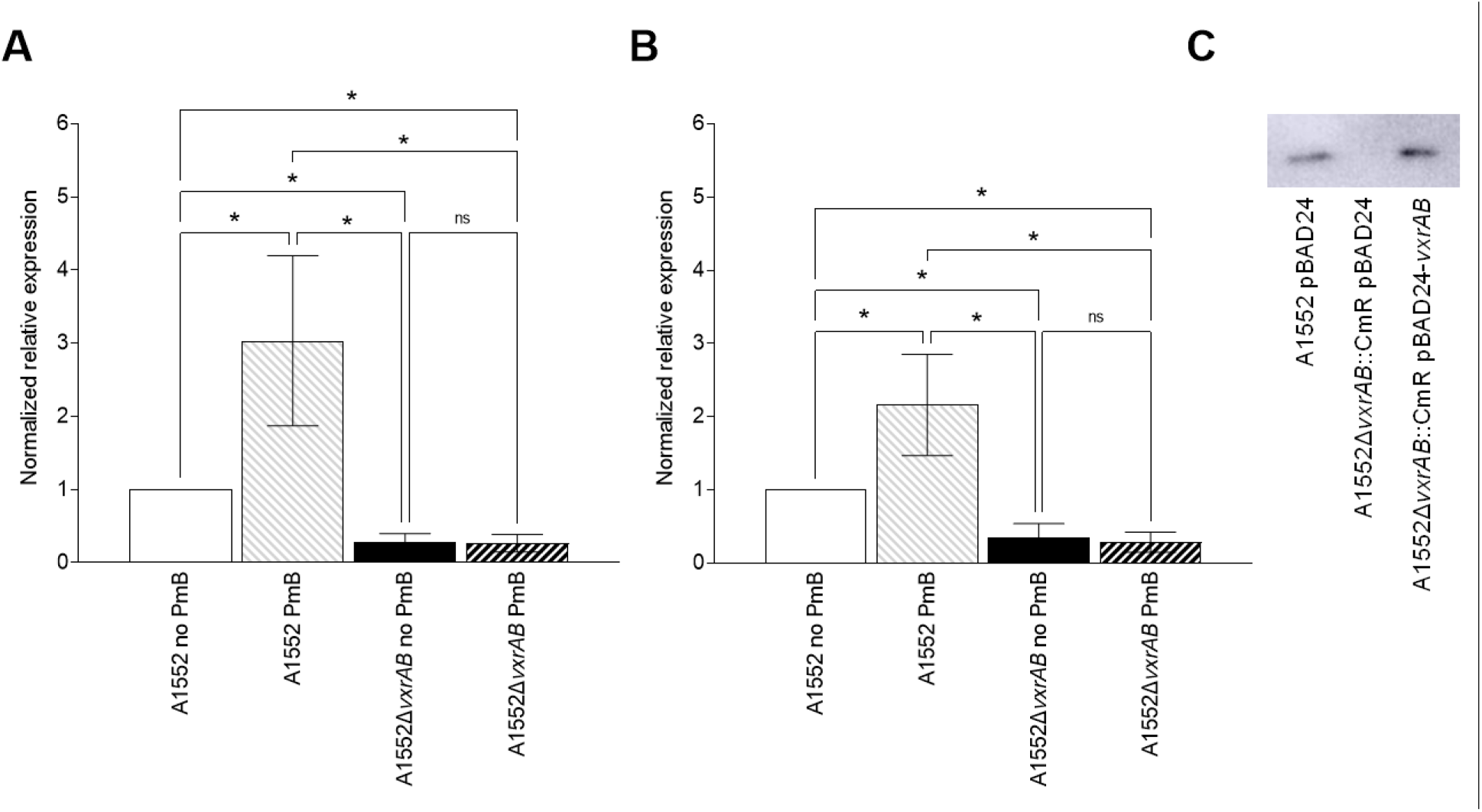
The two-component system VxrAB plays a role in *hcp* upregulation in *V. cholerae* in the presence of subinhibitory concentrations of polymyxin B. *V. cholerae* A1552 and A1552Δ*vxrAB*::CmR (A1552Δ*vxrAB*) were grown to midlog phase in LB2%NaCl with or without 3 μg/ml of polymyxin B (PmB). Total RNA was extracted from cultures and retrotranscribed. The expression of *hcp1* **(A)** or *hcp2* **(B)** was measured by quantitative PCR in the presence of PmB in comparison to the non-treated cells, and normalized using *recA*. Data represent mean ± SD of three independent experiments conducted in triplicates. Asterisk represents a significant difference in expression between treated and non-treated cells (*P*<0.05). ns, non-significative difference. **C)** The complementation of *vxrAB* restores the level of Hcp in the supernatant of A1552Δ*vxrAB*::CmR pBAD24-*vxrAB*. A1552 pBAD24, A1552Δ*vxrAB*::CmR pBAD24 and A1552Δ*vxrAB*::CmR pBAD24-*vxrAB* were grown to late exponential phase in LB-2%NaCl in the presence of PmB.

The secretion of Hcp in A1552 pBAD24, A1552Δ*vxrAB*::CmR pBAD24 and the complemented strain A1552Δ*vxrAB*::CmR pBAD24-*vxrAB* in the presence of PmB was assessed by western blot (Figure 3C). We previously demonstrated that Hcp secretion by *V. cholerae* A1552 is active in the presence of PmB (46). As expected, the secretion of Hcp is greatly reduced upon *vxrAB* mutation (Figure 3C). However, a *vxrAB* complementation completely restored, or even slightly increased, the secretion of Hcp to the level of the WT in the presence of PmB (Figure 3C).

## Discussion

In this study, we aim to determine the impact of subinhibitory concentrations of PmB on the proteome of *V. cholerae* and to identify the regulatory pathways involved in the over expression and over secretion of Hcp in the presence of subinhibitory concentrations of PmB. To do so, we performed a proteomic analysis of the cellular fraction of *V. cholerae* O1 El Tor strain A1552 in the presence and in absence of PmB. We identified a total of 22,819 peptides corresponding to 454 proteins of 7 to 184 kDa, of which 177 had a modified abundance in the presence of PmB.

Several proteins were identified in both our last and current proteomic analysis. It is the case of Hcp that is more abundant in supernatant and bacterial cells in the presence of PmB, and of the proteases VesC and DegP, which was expected since they are secreted components. VesC is secreted through the Type II Secretion System (67), while DegP is secreted by membrane vesicles (68). A putative hydrolase (VC_1485) and an immunogenic protein (VC_0430) were also more abundant in both analysis in the presence of PmB. While the abundance of the lipoprotein Lpp and SucD (α subunit of succinyl-CoA synthetase) was decreased by PmB in the secretome, they were more abundant in the cell fraction. Two proteins were only present in the supernatant in the presence of PmB, and more abundant in the cellular fraction, the amino-acid transporter VC_0010 and the uncharacterized protein VC_0483. One protein, VxrD, was less abundant in both supernatant and bacterial cells in the presence of PmB. Our results show that *V. cholerae* modulates a large proportion of its components in response to PmB, as 39% of the identified proteins had their abundance modified in its presence. Most modulated proteins with increased abundance are implicated in cellular and metabolic processes, with binding, catalytic or transporter activities, that suggests an adaptation to survive the presence of the toxic AMP. This is further supported by the upregulation of efflux systems (VexAB, VprAB, VarF), LPS modifying enzymes (AlmGE), proteases (DegS) and RpoE activating proteins, all known mechanisms of antimicrobials resistance. Numerous proteins displaying a modified abundance in the presence of PmB are yet to be described.

Our previous study has shown that Hcp, the T6SS syringe major component, was more abundant in the secretome of *V. cholerae* under PmB activation (46). The Hcp syringe is wrapped by the contractile sheath composed of multiple helical polymers of VipAB, assembled in an extended form (69–71). This extended form provides enough energy, upon a contraction signal, for a contraction and rotation of VipAB helical polymers that propel the Hcp syringe, along with the effectors, through the bacterial envelope. Based on the fact that, conversely to the Hcp secreted protein, the abundance of VipB, a structural recycled component of the T6SS, was not modified, we hypothesized that PmB might increase the secretion through the T6SS rather than the number of assembled systems at the bacterial surface. Our current proteomic analysis of bacterial cells confirms that the contractile sheath components VipA and VipB are not more abundant in the presence of PmB. ClpV is a AAA+ (ATPases associated with various cellular activities) that hydrolyzes ATP to reshape, or recycle, various substrates (71). In *V. cholerae*, ClpV recognizes the contracted state of the T6SS sheath and forces its disassembly by unfolding VipB (70). To our knowledge, VipA/B are the only recycled proteins through ClpV activity. The recycling of the contractile sheath is essential and facilitates the effectors’ secretion because it makes VipA/B available for further secretion (70, 72). The presence of more ClpV could help the contractile sheath recycling in a cell with a T6SS activated by PmB.

Our proteomic analysis also revealed that the immunity protein TsaB (14, 73), the inner-membrane complex subunit VasK (74), and the FHA protein VasC (TagH), required for secretion (61), are more abundant in the presence of PmB. VasK (TssM) is part of the inner-membrane complex (74) along with VasF (TssL) and VasD (TssJ), a complex randomly distributed in the membrane. Once the inner-membrane complex is formed, it recruits the base plate components, then the Hcp syringe along with the VipA/B sheath (75). It has been suggested that the inner-membrane complexes are pre-assembled in the membrane, more abundant than the syringe portion and that they are reused for further secretion events using new syringe complexes (74). Altogether, our results suggest that more T6SS anchoring complexes might be formed and ready for secretion, while the recycled components abundance remains constant, and the secreted effectors are more produced to increase the number of contraction events and the global T6SS activity in the presence of PmB.

Our proteomic analysis identified many known regulators of the T6SS in *V. cholerae, i.e*. CytR, CRP, TfoY, RpoN, TsrA, OscR and VxrB, but only VxrB had a modified abundance in the presence of PmB. The expression of *crp*, *tfoY, tsrA* and *oscR*, although significantly higher, was only slightly modified by PmB, while the expression of *cytR* and *rpoN* was similar to the control. OscR is a transcriptional regulator that represses T6SS genes expression at low osmolarity (21). CytR is activated upon nucleosides starvation, that further activates the expression of T6SS (23). RpoN, along with the internal regulator VasH, coordinates the expression of the different T6SS genes clusters (6). TsrA is a global regulator (76), while TfoY and CRP are produced in response to c-di-GMP and cyclic-AMP, respectively. While CspV was not identified in our proteomic analysis, our transcriptomic results showed that PmB decreases the expression of *cspV*. However, since its expression is reduced in the presence of PmB and known to be produced upon cold shock (29), it is not surprising that it was not identified in the proteomic analysis. We constructed a mutant in which *cspV* is deleted and compared the expression of *hcp1* and *hcp2* in absence and in the presence of PmB for the wild-type strain and the *cspV* mutant. The expression pattern of both *hcp* genes was similar for the two strains, thus suggesting that *cspV* is not responsible for the increased *hcp* expression in the presence of PmB.

Our proteomic and transcriptomic analysis showed that the presence of PmB increased the production and expression of VxrB, the response regulator of the two-component system VxrAB (34). VxrB is a known regulator of *V. cholerae* T6SS, as its deletion decreases the expression of many T6SS related genes, including Hcp (34). In this study, we constructed a A1552Δ*vxrAB*::CmR mutant, the only essential components of the *vxrABCDE* locus, and a complemented strain. As previously reported, the deletion of *vxrAB* strongly diminished *hcp* expression (34). The complementation of *vxrAB* using pBAD24 restored the secretion of Hcp in the presence of PmB. Conversely to the wild type strain, the presence of PmB did not increase *hcp* expression in A1552Δ*vxrAB*::CmR, demonstrating that it is implicated in the upregulation of *hcp* in the presence of PmB. VxrAB (also known as WigKR) was first identified for its implication in colonization in the infant mouse model and T6SS regulation (34). VxrAB has already been described to regulate other systems implicated in antimicrobial resistance, such as biofilm formation, motility and cell shape maintenance and homeostasis in the presence of cell wall targeting antimicrobials (30, 66, 77). In the later, they demonstrated that the expression of *vxrAB* is increased by antibiotic-induced cell wall damage, and that VxrAB regulates the entire cell wall synthesis pathway, leading to *V. cholerae’s* tolerance to antibiotics (66). Even though we detected no major damages in the cell wall of *V. cholerae* A1552 at a concentration of PmB as high as 25 μg/mL (47), we observed, in this study, a significant upregulation of *vxrB* in the presence of 3 μg/mL of PmB. Taken together, our results suggest that in *V. cholerae* O1 El Tor the T6SS is up regulated by PmB, and that the upregulation involves the TCS VxrAB. The over production of ClpV and VasK also suggests that the T6SS are more efficiently recycled to facilitate the secretion of effectors.

## Acknowledgment

The authors would like to thank Dre Wai from Laboratory for Molecular Infection Medicine Sweden (MIMS) at Umeå University for bacterial strains and the RAQ (Ressources Aquatiques Québec), an inter-institutional group supported financially by the Fonds de recherche du Québec – Nature et technologies (FRQNT) for their support. The proteomic analysis was performed by the Center for Advanced Proteomics Analyses, a Node of the Canadian Genomic Innovation Network supported by the Canadian Government through Genome Canada. This work was supported by the Natural Sciences and Engineering Research Council of Canada (NSERC; http://www.nserc-crsng.gc.ca/index_eng.asp) Discovery grant number RGPIN-2017-05322. AM-D received financial support from the NSERC scholarship program (BESC D3 – 558624 – 2021).

